# Perceptual reorganisation from prior knowledge emerges late in childhood

**DOI:** 10.1101/2022.11.21.517321

**Authors:** G. A. Milne, M. Lisi, A. McLean, R. Zheng, I.I.A. Groen, T. M. Dekker

## Abstract

Perception in the mature human visual system relies heavily on prior knowledge. Here we show for the first time that prior-knowledge-induced reshaping of visual perception emerges gradually, and late in childhood. To isolate the effects of prior knowledge on vision, we presented 4-to-12-year-olds and adults with two-tone images, which are degraded photos that are hard to recognise on first viewing. In adults, seeing the original photo causes a perceptual reorganisation leading to sudden, mandatory recognition of the two-tone version - a well-documented process relying on top-down signalling from higher-order brain areas to early visual cortex. We find that children younger than 7 to 9 years, however, do not experience this knowledge-guided shift, despite viewing the original photo immediately before each two-tone. To assess potential computations underlying this development we compared human performance to three state-of-the-art neural networks with varying architectures. We found that the best-performing architecture behaved much like 4- to 5-year-old humans, who display a feature-based rather than holistic processing strategy akin to neural networks. Our results reveal a striking age-related shift in the reconciliation of prior knowledge with sensory input, which may underpin the development of many perceptual abilities.

## Introduction

The mature human visual system is unique in its ability to accurately recognise objects across a wide range of viewing circumstances. Reaching this level of robustness in object recognition poses a major challenge to computer vision algorithms (Geirhos et al., 2020; Huber et al., 2023; Lindsay, 2021; Pei et al., 2021) and to developing humans. Though infants’ perception of simple shapes is invariant to orientation soon after birth (Slater and Morison, 1987), the ability to recognise and classify more complex objects develops gradually after the first few months of life (Bomba and Siqueland, 1983; Quinn, 2002). This ability likely requires extended visual exploration of objects (Clerkin et al., 2017), and incorporates semantic labelling of visual categories as infants begin to learn language (Smith, 2009; Althaus & Westermann, 2016). By 1-2 years old, infants demonstrate an understanding of the relationship between the visual and lexical categories of familiar objects (Arias-Trejo & Plunkett, 2009), however recognition of these objects under challenging viewing conditions, such as clutter, noise, abstraction, and unusual lighting or orientations, does not become adult-like until at least 10 years old (Bova et al., 2007; Dekker et al., 2011; Nishimura et al., 2009). Current predictive coding models of human vision assign a crucial role to prior knowledge, delivered via top-down pathways in the brain, for parsing ambiguous or cluttered images (Bar, 2004; Friston, 2010; Kersten et al., 2004; Seijdel et al., 2021). Meanwhile, structural and functional MRI measures of long-range neural connectivity that may mediate these feedback signals have been shown to increase continuously over the first decade of life (Baum et al., 2020; Fair et al., 2007). We therefore predict that effective perception of hard-to-recognise objects may develop gradually as long-range neural connections are established across childhood, improving knowledge-based parsing. To track this perceptual development, we asked children aged four to twelve years old to identify ‘two-tone’ image stimuli, which allow us to disentangle the presence of object knowledge from the ability to use this knowledge to inform perceptual inference.

Following their conception by artist Giorgio Kienerk (1869 - 1948) and introduction to psychology by Craig Mooney (1957), two-tone images have offered a famous example of the importance of prior knowledge for recognition. By subjecting greyscale images to Gaussian smoothing and binarisation, visual information is reduced and object boundaries are ambiguated in the resulting black-and-white two-tone (Moore and Cavanagh, 1998). These images are not easily recognisable when viewed naively, but have the intriguing property that once additional cues are made available, recognition becomes near-mandatory in subsequent viewings. This process, referred to as perceptual reorganisation, is thought to involve feedback to low-level visual cortex from higher-order brain areas, including the lateral occipital and prefrontal cortex (Bona et al., 2016; Flounders et al., 2019; González-García et al., 2018; Hardstone et al., 2021; Hsieh et al., 2010; Imamoglu et al., 2012). Pharmacological interference of top-down pathways reduces the similarity of responses in primary visual cortex (V1) to two-tone and greyscale images, suggesting a causal role of these feedback connections in perceptual reorganisation (van Loon et al., 2016). This top-down information processing may operate by enhancing visual sensitivity to low-level image features of the two-tones computed in V1, such as orientation and edge information (Teufel et al., 2018). This is consistent with Bayesian models of perceptual inference, where priors are propagated to lower areas via top-down processes and combined with bottom-up information to shape perception (Kersten et al., 2004; Lee and Mumford, 2003).

Two-tone image recognition thus provides a well-controlled paradigm for studying how the use of prior knowledge to parse ambiguous images develops. Though Mooney himself already showed that the ability to naively recognise two-tones of faces is impaired during childhood and improves with age (Mooney, 1957), less is known about children’s ability to recognise non-facial two-tone stimuli, and even less so about the development of perceptual reorganisation. In their book chapter on binocular perception, Kovács and Eisenberg (2005) hypothesised this ability may also be impaired in young children. In a 2007 conference paper, Yoon and colleagues then reported low accuracy locating object features on two-tones in a small sample of three- to four-year-olds, despite them having seen the original photo, in line with reduced perceptual reorganisation (Yoon et al., 2007). Thus, across childhood there may be drastic changes in how prior knowledge guides perceptual inference about ambiguous images, which may have broad-reaching implications for the development of object recognition. However, this is currently underappreciated in the literature, and a characterisation of two-tone image perception across childhood that systematically tests this is currently lacking.

Here we address this gap by testing perceptual reorganisation and two-tone image processing throughout childhood in 72 four- to twelve-year-olds. Crucially, we use methodological and analytical approaches that systematically account for potential confounding factors besides the ability to use prior knowledge to parse two-tones that also change with age. These include object knowledge, task comprehension, and response execution. We explore which computational mechanisms may underlie shifts from child-like toward adult-like two-tone recognition by investigating image parsing and feature extraction strategies used at different ages. We compare human performance to that of Convolutional Neural Networks (CNNs) to investigate the extent to which the architectural properties of these models, i.e., incorporation of feedback processing or computational depth, can emulate early stages of human visual development. We find striking differences in the way that young children, and CNNs, parse ambiguous two-tone images, whereby adult levels of prior-knowledge-driven perception do not develop until late childhood. As in computational object classifiers, we link children’s reduced recognition of two-tone images to a more featural-based processing strategy.

## Results

Four- to five-year-olds (n = 31), seven- to nine-year-olds (n = 23), ten- to twelve-year-olds (n = 18) and adults (n = 13) sequentially viewed two-tone images on a touchscreen before and after cueing with the corresponding greyscale image. In each trial, participants were asked to name the content of the naively-viewed two-tone (Figure 1A, stage 1), after which the image transitioned into the corresponding greyscale to cue object knowledge, and participants were again asked to name the content to confirm they could recognise the image (Figure 1A, stage 2). To measure perceptual reorganisation, this image transitioned back to the same two-tone and participants were asked to touch the locations of two characteristic object features (Figure 1A, stage 3). After completing trials for twenty two-tones and five Catch images (unsmoothed two-tones), participants were shown each greyscale image again and asked to touch the same target features that were prompted for the corresponding two-tone images (Figure 1A, stage 4).

**Figure 1:**
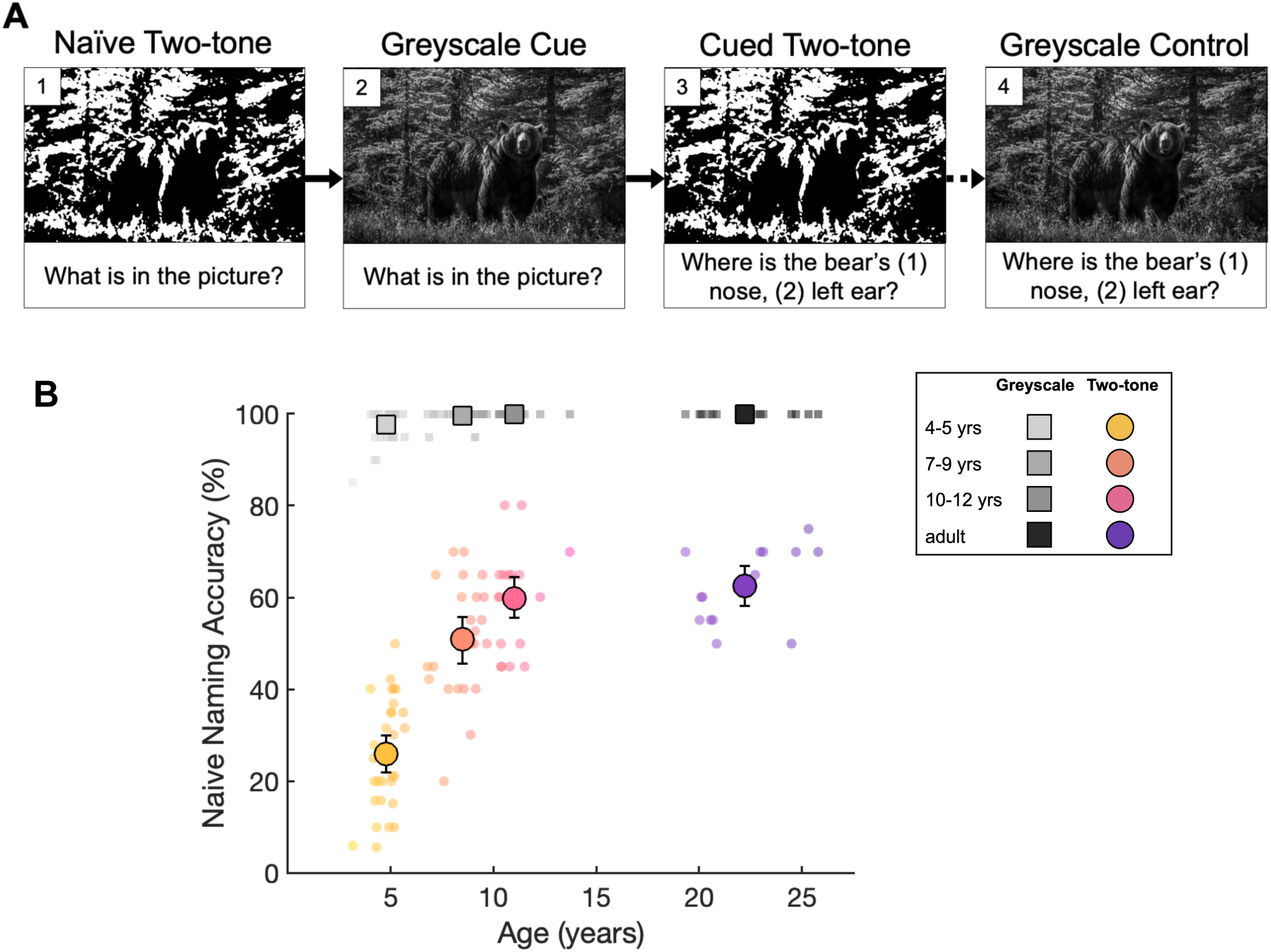
Experimental trial schema and Naive two-tone recognition accuracy **A)** An example trial for one image, stages 1 (naive two-tone naming), 2 (greyscale cueing and naming), and 3 (cued two-tone pointing) are sequential, and stage 4 (greyscale control pointing) occurs separately after the main task. **B)** Naming accuracy of Naive two-tones (coloured circles; see stage 1 in A), and greyscale images (grey squares; see stage 2 in A). Small markers show participant means, large markers show age group means, error bars show bootstrapped 95% Confidence Intervals.

### Accounting for age differences in greyscale recognition

To account for potential age differences in object knowledge, we first established that greyscale naming accuracy was high and matched at all ages (Figure 1B, grey squares, Χ^2^(3) = 3.58, *p* = 0.31). Nevertheless, trials in which greyscales were incorrectly named were excluded from all further analyses (total trials excluded for 4- to 5-year-olds = 15, 7- to 9-year-olds = 2, 10- to 12-year-olds and adults = 0).

### Age differences in Naive two-tone recognition

In contrast to the accurate naming of the greyscale images at all ages, a logistic mixed-effects model (see Methods for details) revealed that naive naming accuracy for the two-tone versions of these greyscales was lower, and improved substantially with age (Χ^2^(3)=89.88, *p*=2.2e-16; Figure 1B, coloured circles). Wald tests on model coefficients revealed that 4- to 5-year-olds (*z* = −9.6; *p* < 2.0e-16) and 7- to 9-year-olds (*z* = −3.2; *p* < 1.38e-3) were less accurate than adults, while 10- to 12-year-olds performed similarly to adults (*z* = −0.70, *p* = 0.48). This age difference is unlikely due to failure to comprehend or comply with the task since performance for easily recognisable Catch images was high for all ages (see Supplementary Figure 2A). Mean naming accuracy per image for naively viewed two-tones was correlated between all age groups (r_pearson’s_ > 0.7, *p* < 5.71e-4, see Supplementary Figure 4 for all correlations). Together this suggests that up to at least 7 to 9 years of age, recognition of familiar objects is more impaired by two-tone transformation than in adulthood, but that the features that make a two-tone hard to recognise at first sight remain qualitatively consistent across age.

### Age differences in perceptual re-organisation

Perceptual reorganisation was assessed by measuring two-tone recognition immediately after presenting participants with the greyscale cue. To minimise memory load and make clear that greyscales and two-tones were different versions of the same image, all images were ‘morphed’ into one another by fading from the greyscale to the overlaid two-tone with position and dimensions maintained. Participants were then asked to touch two predetermined features on each two-tone on the touchscreen (Figure 1A, stage 3). To ensure participants could correctly recognise and point out all targets, they were also asked to locate the corresponding features on the original greyscale images in a control task after the main experiment (Figure 1A, stage 4). Touched coordinates were scored by two methods; first, the percentage of touched points falling within researcher-defined regions gives ‘pointing accuracy’, and second, the distance between corresponding touched coordinates on two-tone and greyscale images gives ‘pointing distance’ (see Figure 2A).

**Figure 2:**
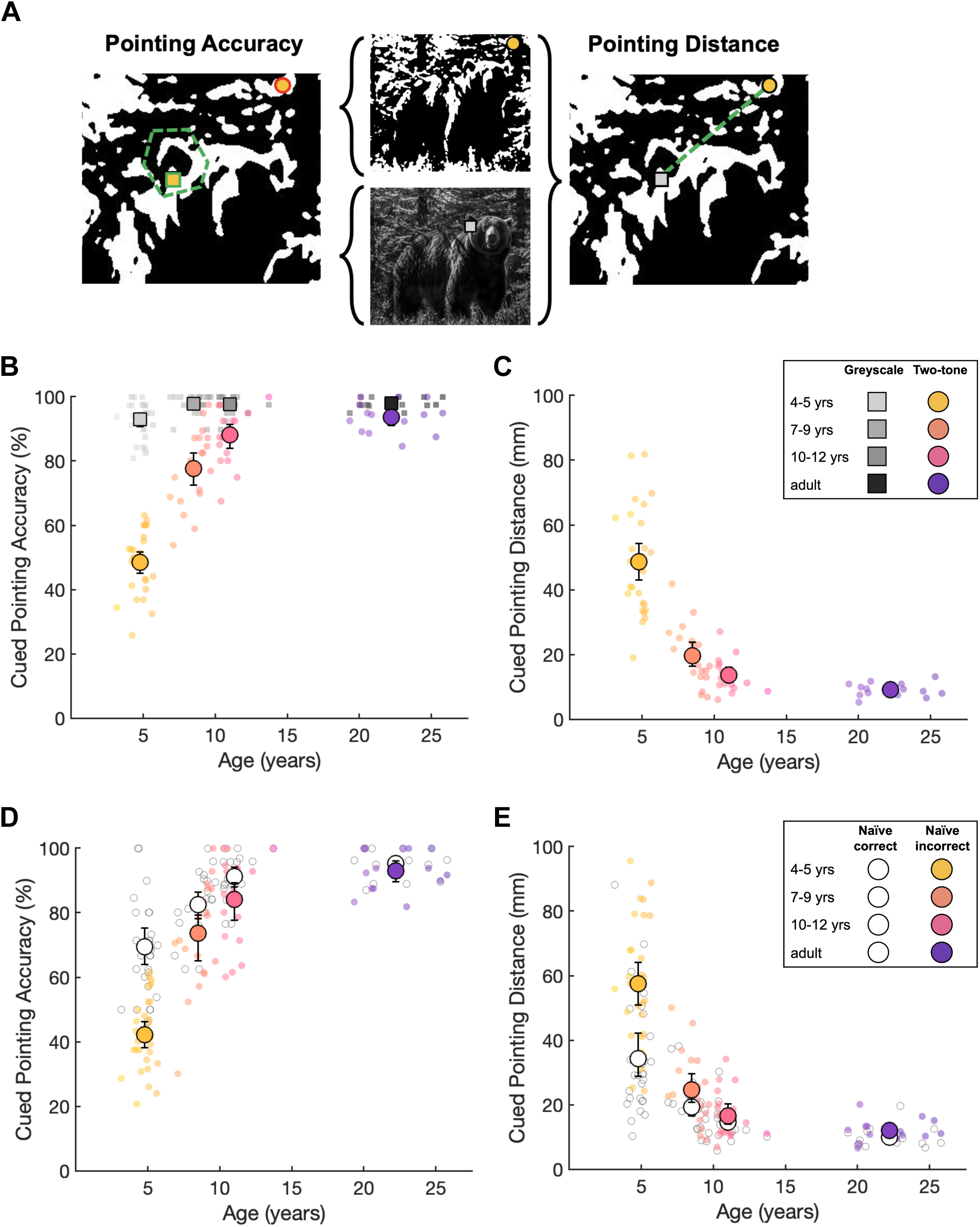
Cued two-tone recognition measured by Pointing Accuracy and Distance **A)** Circular and square markers show example two-tone and greyscale cued pointing responses for one of two targets respectively. Touched points were scored via two methods; the dashed polygon shows the example ‘correct’ feature location used to determine ‘pointing accuracy’ (green outlined marker scored as correct and red outlined marker scored as incorrect). The dashed line represents the distance between touched points for the corresponding target in the two-tone and greyscale conditions, defining the ‘pointing distance’ measure. **B)** Pointing accuracy for two-tones (coloured circles) following greyscale exposure, and greyscales (grey squares), as measured by percentage of touched locations falling within the predefined correct area for each target (Pointing Accuracy). Small markers show participant means, large markers show age group means, error bars show bootstrapped 95% Confidence Intervals. **C)** Pointing Distance between touched locations on Cued two-tone and greyscale conditions of each image in mm (coloured circles; all images were displayed at a fixed height of 273.9 mm). Markers and error bars are as in Panel B. **D)** Pointing accuracy of previously recognised (white circles) and previously unrecognised (coloured circles) two-tones following greyscale exposure. Markers and error bars are as in Panel B. **E)** Pointing distance of previously recognised (white circles) and previously unrecognised (coloured circles) two-tones following greyscale exposure. Markers and error bars are as in Panel B.

There were large age-related changes in cued two-tone recognition, with Pointing Accuracy and Distance both improving substantially with age (Figure 2B & 2C, coloured circles). In contrast, for greyscale and Catch images, Pointing Accuracy was high across all ages (Figure 2B, grey squares, and Supplementary Figure 2B, white circles), so this age difference is unlikely to reflect general changes in task comprehension. However, it is possible that these age differences are driven by factors unrelated to perceptual reorganisation, such as adults having recognised more two-tones naively, or children exhibiting greater imprecision or different pointing strategies despite recognising the content. Therefore, to test for perceptual reorganisation whilst accounting for naive two-tone recognition and pointing skills, we compared pointing performance within subjects for trials in which two-tones were naively recognised before cueing (and therefore likely still recognised after) to trials in which they were not. If greyscale cueing effectively induced perceptual re-organisation, pointing performance should be equivalently high across these two trial groups, irrespective of initial recognition. Furthermore, in our mixed-effects modelling analysis (see Methods for details) we accounted for random effects of participants, individual images, ROI size, and unequal numbers of trials.

This analysis showed that a difference in cued pointing performance for naively recognised and unrecognised two-tones was largest for 4- to 5-year-olds and decreased with age (for pointing accuracy: Χ^2^(3)=13.85, *p*<3.11e-3, Figure 2D; for pointing distance: Χ^2^(3)=16.24, *p*=1e-3, Figure 2E), with greater benefits of cueing for older participants on both recognition indices. Wald tests on model coefficients showed that in adults, feature localisation was equivalently accurate for naively recognised and unrecognised two-tones (pointing accuracy: *z*=-1.24; *p*=0.21; pointing distance: *z*=0.25; *p*=0.8), in line with pervasive perceptual reorganisation in the mature system. For pointing accuracy, the greyscale cue was significantly less beneficial for 4- to 5-year-olds than for adults (*z*=3.22, *p*=1.26e-3), marginally less beneficial for 7- to 9-year-olds (*z*=1.71, *p*=0.08), and adult-like for 10- to 12-year-olds (z=1.46, *p*=0.14). For pointing distance, no pairwise age group comparisons reached statistical significance, potentially because this measure was more variable. Image difficulty (number of errors for naming accuracy or pointing accuracy per image) was significantly correlated across naive and cued conditions for 4- to 5-year-olds only (r_pearson’s_= 0.50, *p* = 0.02; *p* > 0.09 for all other groups, see Supplementary Figure 4 for all correlations). Together, these data reveal that processes supporting adult-like perceptual reorganisation, in which two-tone parsing is qualitatively altered when prior knowledge is made available, develop gradually over the first 10 years of life.

### Age differences in cued two-tone parsing strategies

To investigate age differences in image parsing strategies of cued two-tones, we compared cued pointing responses across ages. We have shown above that adults typically located both targets on cued two-tones regardless of naive recognition, whilst 4- to 5-year-olds often failed to locate these targets, and showed poorer performance for naively unrecognised images in particular. Cued pointing patterns reveal that when failing to locate the target feature, 4- to 5-year-olds still seem to display a task-oriented strategy, as illustrated by a high prevalence of local feature-driven errors for this age group (Figure 3). That is, the incorrect selection of two-tone features that canonically resemble the target feature, but whose location does not correspond with the global image content (‘local errors’). For example, when asked to locate features of a woman’s face, many children instead located corresponding features of a pareidolic face appearing on the woman’s forehead, despite performing accurately and precisely on the same task in the greyscale condition (Figure 3Ai). Likewise, when asked to point to the cowboy’s hat and horse’s ear, many erroneously located ‘hat-like’ and ‘ear-like’ shapes, regardless of their distance from the target features in the just-seen greyscale image (Figure 3Aii). Similarly, when the outlines of target features were obscured following two-tone transformation (*e.g.*, the noses of the fox [Figure 3Aiii, Target 1] and panda [Figure 3Aiv, Target 2]) many children incorrectly selected nearby shapes with more defined outlines. To quantify this pattern of responses across age groups, we measured the proportion of errors displaying a locally feature-driven strategy for eight targets for which incorrect features that matched the target’s shape properties were present in the two-tone (Figure 3B). We found that whilst adults did not commit any such errors for these targets, the proportion of errors that could be attributed to a local feature-driven strategy decreases dramatically with age. Together, this suggests that young children who showed little benefit of cueing on image recognition made errors consistent with prioritisation of local features, rather than holistic image processing.

**Figure 3:**
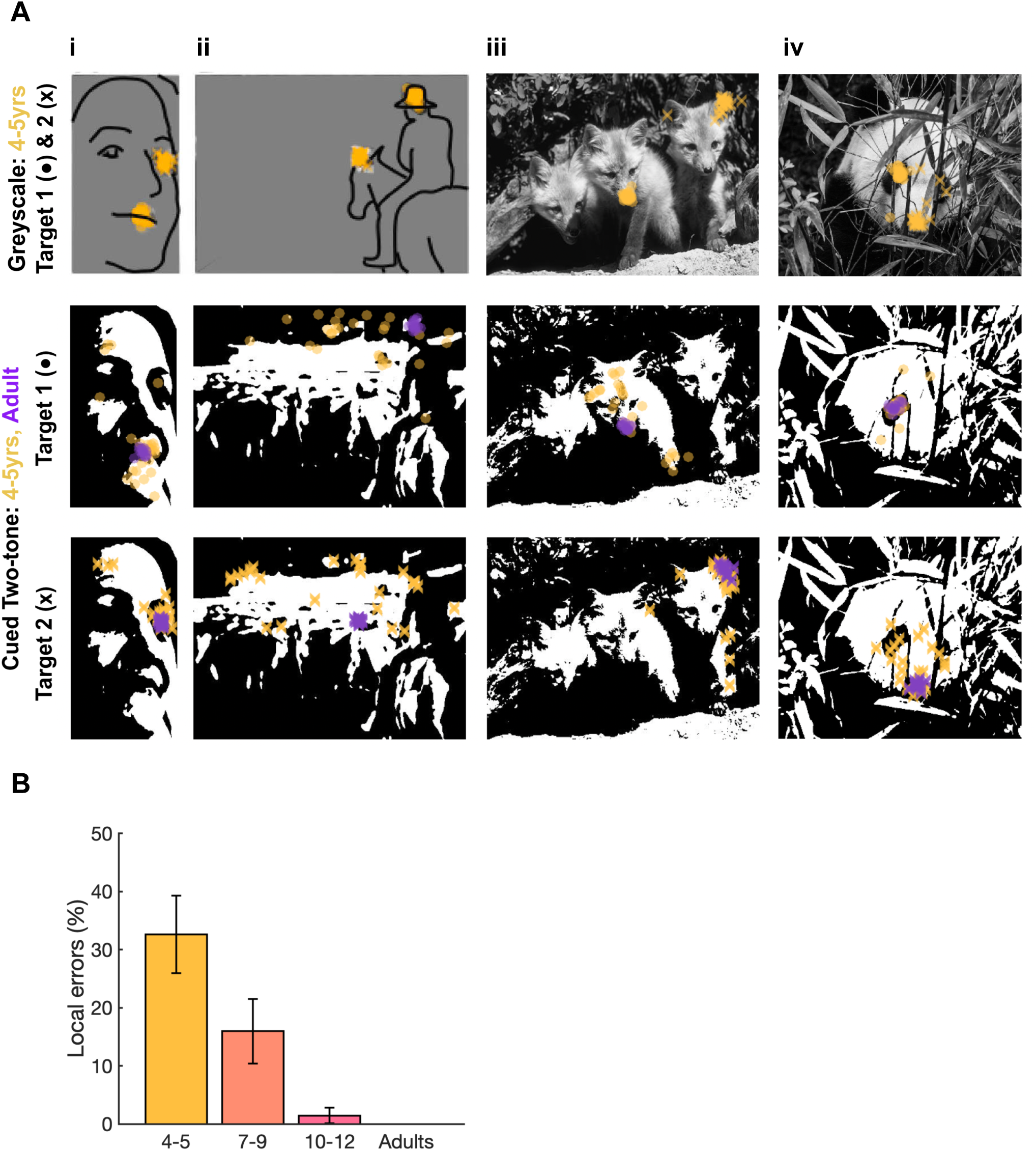
Analysis of Cued Pointing errors **A)** Top row: 4- to 5-year-olds’ greyscale pointing for targets 1 (circles) and 2 (crosses), target prompts were i) ‘the lady’s mouth’ (1) and ‘the lady’s right eye’ (2), ii) ‘the cowboy’s hat’(1) and ‘the horse’s left ear’ (iii), C ‘the middle fox’s nose’ (1) and ‘the right fox’s right ear’ (2), and iv) ‘the panda’s left eye’ (1) and ‘the panda’s nose’ (2). Middle and bottom rows show 4- to 5-year-olds’ (yellow) and adults’ (purple) cued two-tone pointing for targets features 1 and 2 respectively. **B)** Mean proportion of incorrect responses that constituted ‘local errors’ (i.e., the selected feature matches the local shape properties of the target, but is positioned incorrectly with regards to global image content) by age group for 8 targets where such locally-matching features were present. Error bars show standard error.

### Human development versus computational image-recognition models

We next compared human performance on two-tone recognition to that of Convolutional Neural Networks (CNNs) to test whether different architectures may correspond to sequential stages in human development. We selected three image recognition models based on their architectural properties and availability; (1) AlexNet (Krizhevsky et al., 2017) - a shallow feedforward network extensively used in computational neuroscience studies (Lindsay, 2021), (2) CorNet-S, a biologically inspired shallow network that incorporates feedback within late and early-stage processing layers (Kubilius et al., 2019), and (3) NASNet-Large - a deep feedforward network with a complex branching architecture designed by auto machine-learning algorithms optimising for transferable image classification (Zoph et al., 2018). All three models are publicly available and were pre-trained on 1.3 million images from the ImageNet training set into 1000 different classes (Deng et al., 2009).

To assess the possibility of improved two-tone recognition of ‘familiar’ images compared to novel images (akin to perceptual reorganisation) in these CNNs, models were tested on two image sets. First, nineteen of the above-used two-tones that were not included in ImageNet (the data set used to train all three models) formed the ‘novel’ image set, and second, twenty comparable two-tones created from ImageNet images formed the ‘trained’ image set. CNNs with the capacity to benefit from image cueing would be expected to more accurately classify two-tones from ‘trained’ as opposed to ‘novel’ image sets. This, however, was not the case for any of the models tested. It is unlikely that this reflects differences in difficulty levels across the two image sets; image pairs across the sets were matched on low-level statistics, piloting in an additional nine adults ensured that two-tone recognition performance was equivalent across the two sets (novel set: 61% naive accuracy, 94% cued accuracy; trained set: 63% naive, 89% cued accuracy), and all CNNs showed comparable performance on the greyscale images of each set (Figure 4A).

**Figure 4:**
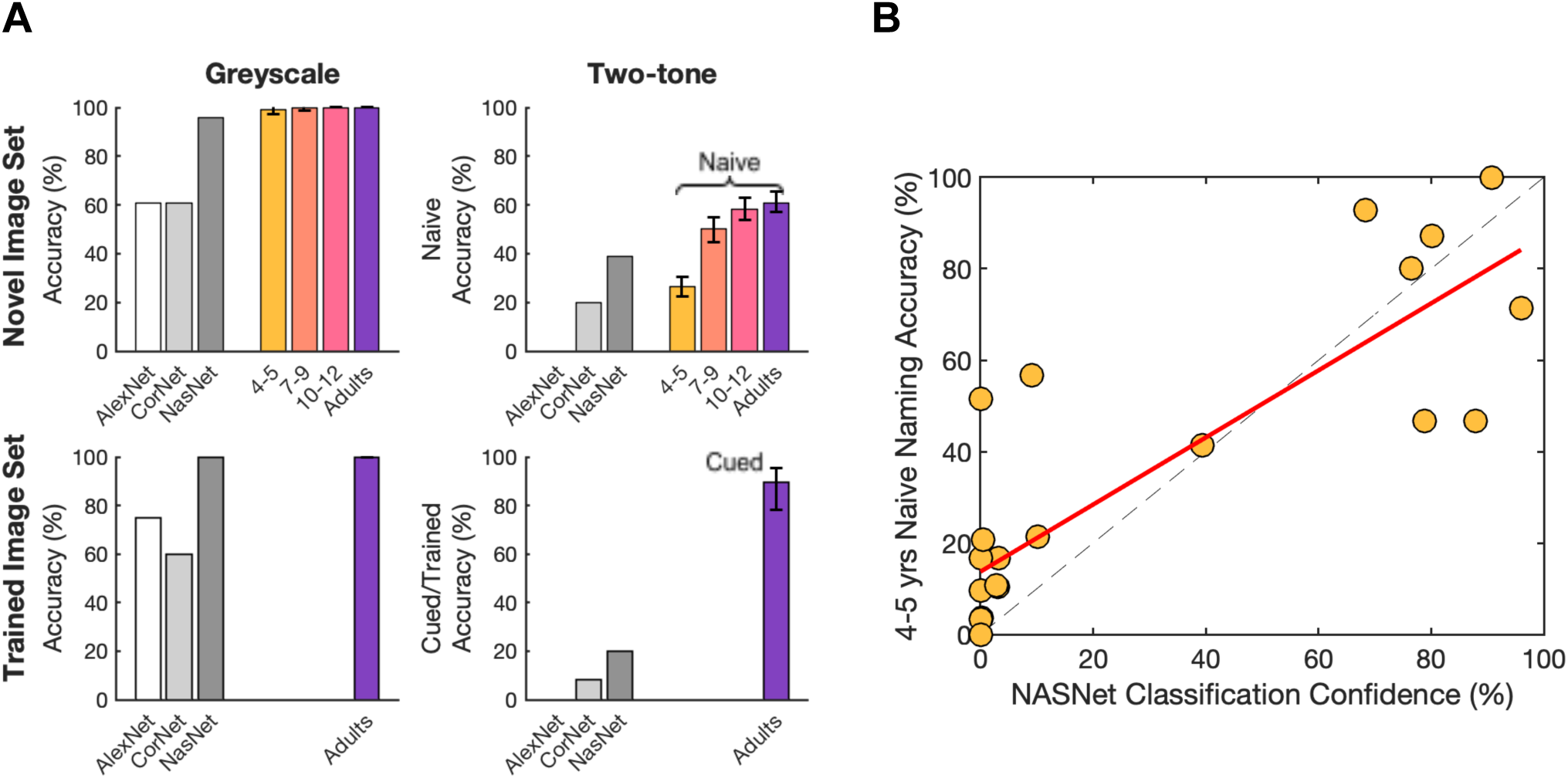
Comparison of CNN performance to human performance **A)** Image recognition accuracy of CNNs (AlexNet, CORnet-S, NASNet-large; greyscale bars) and human performance (coloured bars, error bars show bootstrapped 95% confidence intervals) for each condition (greyscale and two-tone) of images used in current study (Novel Image Set) and Image Net images used to train CNN models (Trained Image Set). CNN performance: percent of images correctly labelled; human performance: Naive Naming Accuracy. **B)** Image-wise comparison of mean 4- to 5-year-old Naive Naming Accuracy and NASNet performance (measured by probability weighting of the highest-scoring correct label) for Catch and two-tone trials of Novel Image Set. Red line shows linear least-squares line of best fit, dashed line shows x = y.

We next tested how CNN performance compared to that of human participants for naive recognition. We found that NASNet-large was the highest-performing model for both greyscale and two-tone images but nevertheless only reached the performance level of the youngest age group for naive two-tone classification. While both CORnet-S and AlexNet had comparatively lower performance for greyscales than NASNet, for two-tones CORnet did approach naive performance levels of 4- to 5-year-olds whilst AlexNet failed to classify any two-tones correctly (Figure 4A). Interestingly, NASNet’s classification confidence for two-tones from the ‘novel’ image set, quantified as the probability weighting of the highest-scoring correct label, correlated across images with human naive naming performance for all ages (Figure 4B, plotted for 4- to 5-year-olds, r_pearson’s_ = 0.84, *p* = 4.98e-7; see Supplementary Figure 4 for all correlations). This suggests that despite vastly different computational implementations, there is a common aspect of these two-tone stimuli that poses a challenge to both human vision and CNNs, which adults can overcome when object knowledge is made available.

It is well-known that CNNs are vulnerable to image manipulations and have difficulty generalising to out-of-distribution datasets (Geirhos et al., 2020). To explore which factors in particular make two-tone images hard to recognize for developing humans and CNNs, we assessed correlations of two-tone recognition with various low-level image features and parameters of the two-tone transformation. Given the proposed need for prior knowledge to resolve edge information for two-tone recognition (Teufel et al., 2018; Moore and Cavanagh, 1998), we assessed how two-tone transformation altered various summary statistics of image edge information. Firstly, edge density, as measured by a Sobel edge detector (Mathworks, 2020) was reduced by two-tone transformation as expected (Supplementary Figure 5B), and larger reductions were associated with poorer naive two-tone recognition in 4- to 5-year-olds and NASNet (r_pearson’s_ > 0.5 and *p* < 0.01 for both). Next, we considered two biologically principled measures of scene statistics, Contrast Energy (CE) and Spatial Coherence (SC), computed via spatial image filters that emulate parvo- and magnocellular operations respectively (Groen et al., 2013; see Supplementary Materials 5 for details). Two-tone transformation was found to increase CE but decrease SC, with greater alterations corresponding with poorer two-tone recognition for 4- to 5-year-olds (see Supplementary Table 5). Together, these correlations may suggest that object recognition in younger children is more sensitive to disruption of edge information in the image than adults, but note that these analyses may be limited by the narrow score ranges in older age groups. However, the extent to which these image statistics correlated with performance did not exceed that of the smoothing levels applied during the two-tone transformation. As expected, increased smoothing was associated with poorer naive two-tone recognition for all ages (r_pearson’s_ < −0.6 and *p* < 0.001). It was also associated with poorer naive two-tone classification for NASNet and cued two-tone performance for children below the age of ten (see Supplementary Table 5). These results suggest that the tested measures of scene statistics do not explain what contributes to two-tone difficulty beyond the amount of information that is lost via smoothing.

## Discussion

To test how prior knowledge guides visual object perception between the ages of 4 and 12 years, we sequentially presented ambiguous two-tone images and greyscale cues. In adults, information about a two-tone’s content provided by the original greyscale induces perceptual reorganisation, resulting in near-mandatory recognition of previously unrecognisable stimuli. To quantify how this process develops across childhood, we compared the ability to locate features on two-tones that were not recognised naively (i.e., candidate images for perceptual reorganisation) to the same measure for two-tones recognised on first viewing. Here, performance on these naively recognised two-tones provides an individual benchmark for feature localisation error when image content is perceived. So, if greyscale cueing causes accurate recognition of previously unrecognised two-tones (perceptual reorganisation), pointing performance should approach this benchmark. Crucially, this comparison isolates the effects of prior knowledge on two-tone recognition, as age differences in other task-relevant abilities (e.g., pointing precision, biases) should affect both conditions equally.

We first showed that naive two-tone recognition underwent substantial developmental improvement, increasing from 26% to 63% correct between the ages of 4-5 years and adulthood. However, despite this large difference in the effect of visual information loss in two-tones, correlations between image difficulties across all age groups showed that the processes hindering naive two-tone recognition remain consistent across age. After seeing the original greyscale, adults and the oldest children could locate target features on previously unrecognised two-tones with similar accuracy as for the two-tones they could recognise naively, clearly demonstrating perceptual reorganisation. However, younger children experienced drastically less new recognition of two-tones after seeing the greyscale cue, pointing out substantially fewer image features correctly than their benchmark for naively recognised two-tones. In addition, image difficulty before and after greyscale cueing (i.e., imagewise naive naming and cued pointing accuracy) was correlated for young children, but not for older children and adults. This indicates that only at older ages does prior knowledge induce a qualitative shift in how two-tones are processed. Together, these results reveal that the effective use of prior knowledge in perception poses a major challenge to human development, with improvements occurring gradually between the ages of 4 to 12 years.

It is highly unlikely that this developmental shift in perceptual reorganisation reflects age differences in object knowledge, task comprehension or non-perceptual processes. While the two-tone paradigm intrinsically minimises potential differences in participants’ object knowledge by providing the relevant information in a task-relevant format, we further controlled for object knowledge differences by only including trials in which greyscale images were correctly named. Task comprehension was also ensured for all participants, both during an initial training phase and throughout the task, via ‘easy’ two-tone images (Catch images, made by thresholding greyscale images without applying smoothing). High recognition performance on these trials (Supplementary Figure 2) demonstrates that all age groups understood, remembered, and followed instructions for both two-tone recognition tasks (naming and pointing) throughout the experiment. In addition, the use of visual stimuli and blurred transitions reduces the demand for working memory, cross-modal integration and configurement of mental representations. Alternative cueing methods, such as non-sequential paradigms and semantic cues, are sufficient to trigger perceptual reorganisation in adults (González-García et al., 2018; Ludmer et al., 2011; Nordhjem et al., 2015; Samaha et al., 2018). While in the present study we prioritised reducing load for cross-modal and working-memory processes known to concurrently develop throughout childhood (Buss et al., 2018; Jüttner et al., 2006; Swanson, 2017), future work should test how these processes affect cue integration across development.

To explore what perceptual strategies younger children employed to parse cued two-tones, we reviewed the patterns of locations that 4- to 5-year-olds selected on these images. This revealed that children often located correct object features if these were clearly outlined, but were prone to large localisation errors when object feature boundaries were obscured or disrupted, instead selecting salient features that plausibly matched the target in outline (*e.g.*, when asked to locate a hat, many identified a shape with a hat-like outline, albeit in the wrong location). This error pattern suggests that younger children looked for features by relying on local shape outlines, rather than by identifying and grouping shapes relevant to the object gestalt. Similar strategies have been reported in adults with Autism Spectrum Disorder, who have outperformed non-autistic adults in tasks that favoured perception of local rather than global features (Happé and Booth, 2008; though see Van der Hallen et al., 2015). These individuals also show evidence of reduced effects of prior knowledge of two-tone processing (Król and Król, 2019; though this may be specific to face stimuli: Loth et al., 2010; Van de Cruys et al., 2018), which has been attributed to a relative downweighting of prior knowledge with respect to sensory inputs. Differences in weighting sensory input and prior information have also been implicated in numerous neuropsychiatric disorders (Fletcher and Frith, 2009; Park et al., 2022; Shanmugan et al., 2016). Indeed, populations that suffer from hallucinations and psychosis-prone individuals experience higher levels of perceptual reorganisation of two-tones than healthy adults (Davies et al., 2018; Teufel et al., 2015; Zarkali et al., 2019), attributed to an upweighting of prior information to compensate for increased noise levels in bottom-up sensory streams (Davies et al., 2018; Fletcher and Frith, 2009; Kapur, 2003; Rivolta et al., 2014; Teufel et al., 2015). The severity of symptoms associated with these conditions, and the extensive research ongoing to alleviate them, reemphasizes the importance of characterising the normal development of top-down processes. Further, many of these disorders are diagnosed in late childhood or adolescence, concurrent with the dramatic changes in top-down weighting shown here. It is therefore an important possibility that the developmental shift we report here could be involved in the onset of neurological pathologies.

Like the youngest children in this study, Convolutional Neural Networks (CNNs) have also been shown to prioritise local over global information for object recognition, which too has been linked to higher levels of dependency on feedforward information processing when compared to the adult human visual system (Baker et al., 2018; Brendel and Bethge, 2019; Geirhos et al., 2019). To assess if this, or other image-processing differences of CNNs, correspond to distinct levels of two-tone recognition, we compared human two-tone recognition to that of CNNs with distinct network architectures. None of the models tested reached naive adult performance, or showed any recognition improvements for ‘familiar’ two-tones (made from the image set on which models were trained (Deng et al., 2009)) compared to novel two-tones, despite matching for difficulty. However, NASNet-large (Zoph et al., 2018), the deepest model tested and only one to achieve human-like greyscale recognition, reached the two-tone performance level of 4- to 5-year-old children. Whilst CORnet (Kubilius et al., 2019) and AlexNet (Krizhevsky et al., 2017) showed similarly poor levels of greyscale recognition, CORnet performed better on two-tones for both trained and novel image sets. These results suggest that training with the original image used to make a two-tone may not be sufficient to trigger an analogue of perceptual reorganisation in these CNNs. Due to the common training set of these models, the differences in naive two-tone recognition seen here are likely driven by architectural features. Namely, increased model depth may have afforded NASNet-large improved performance for both greyscale and two-tone images compared to the other shallower models, whilst the addition of recurrency may drive CORNet’s advantage for two-tones over AlexNet, which is both shallow and completely feedforward. It is important to note, however, that it is possible for purely feedforward models to achieve equivalently ‘recurrent’ computations as CNNs with feedback connections (van Bergen and Kriegeskorte, 2020), which may explain why NASNet-large was able to outperform a more biologically-inspired model. Of course, CNNs differ greatly from the human visual system; in addition to the local biases discussed above, Geirhos and colleagues (2019) found that CNNs prefer images that are coloured or high-contrast. These biases may have inhibited general recognition of our stimuli set, none of which included colour information, but less so for Catch Images, which were high-contrast but unsmoothed (Supplementary Figure 2C). Interestingly, despite large mechanistic differences, NASNet and human participants showed correlated imagewise performance on naive two-tones, revealing commonalities in the challenges these images pose for machine and human visual systems, especially those still developing.

To explore what these challenges may be, we tested how low-level spatial image properties correlated with two-tone recognition performance in developing humans and CNNs. As expected, the smoothing and thresholding levels used in the two-tone transformation process negatively correlated with naive recognition across ages in humans, and for NASNet. This is unsurprising, as higher smoothing levels result in a greater loss of information, and higher thresholds effectively increase image shadows. As these image manipulations can also be expected to interact in a non-linear or unpredictable manner, we used physiologically plausible models of image statistics extraction to quantify the effects of two-tone transformation on low-level spatial image properties (i.e., edge energy, contrast energy, and spatial coherence) computed at early visual processing stages (Groen et al., 2013). While the effects of smoothing and binarising images clearly shifted the low-level spatial properties, we did not find that these statistics correlated better with performance than smoothing levels alone. Nonetheless, as only 24 images were included to ensure a child-friendly task duration, this study was not optimised to model the effect of image properties on perception. Studies with larger image sets that can systematically compare the effects of different image features will therefore be necessary to assess the bottom-up contributions to two-tone perception across development.

Another factor that may contribute to the development of two-tone perception is the efficiency of spatial integration mechanisms across the cortical hierarchy. Two-tone perception requires both the detection and segmentation of object contours from irrelevant contours (i.e., from cast shadows, occluders and background objects) and the integration of these relevant contours to form the figural percept (Moore and Cavanagh, 1998; Poltoratski and Tong, 2020). There is evidence that contour integration develops substantially in childhood: Kovács et al. (1999) showed that the ability to detect contours consisting of collinear-oriented elements amidst misaligned distractor elements improves substantially between 4-12 years of age. Unlike adults, children were less tolerant to distractors when there were larger spacings between the contour elements, suggesting a limitation on spatial integration distance rather than a reduced ability to detect signal in noise. Similarly, the perception of illusory ‘Kanizsa’ shapes, in which shape corners are visible but the contours connecting these corners are not, has been shown to develop gradually over the first decade of life (Nayar et al., 2015). Though these stimuli are simpler than two-tones, the perception of Kanizsa shapes has also been shown to involve hierarchically-organised feedback processing (Kok et al., 2016; Wokke et al., 2013). More complex images and those with less clearly defined object boundaries have been shown to require higher levels of recurrent processing in order to extract the image content (Groen et al., 2018; Kirchberger et al., 2021); a process that can be silenced by disrupting higher-order visual areas (Kirchberger et al., 2021; Wokke et al., 2012). An increase in top-down signal integration with sensory inputs across childhood may therefore provide a common explanation for the prolonged development of these perceptual tasks.

While we carefully matched visual object knowledge of the two-tone across age groups, it is possible that the primed object representation was still less abstract, or ‘invariant’ to image distortions, in children (and CNNs), thus offering a less robust template for parsing the two-tone. Indeed, in a cross-cultural study, adults from an isolated tribe with little-to-no experience with pictorial representations benefitted less from greyscale cueing than Western adults (Yoon et al., 2014). We made every effort to make clear that two-tones and greyscales were two versions of the same photos by blurring corresponding images into each other. It is however possible that benefiting from greyscale cueing requires an understanding of dual representations, which children have been shown to lack (DeLoache et al., 1997). It thus remains to be tested whether children (or computer vision models) would experience higher levels of perceptual reorganisation after extensive training with the depicted objects in varying orientations, sizes, and depictions, and/or the two-tone form.

In sum, we show that throughout most of childhood the benefit of prior knowledge on two-tone recognition is drastically reduced compared to in the mature visual system. When compared to adults, young children may focus more heavily on local image features, characteristic of bottom-up processing streams. We found qualitative evidence of these strategies in young children, whose image recognition also correlated with the performance of a CNN - neural networks that have previously been shown to exhibit local biases in image processing. For both these visual systems, the development of prior-knowledge-driven perception, a central aspect of adult human vision, could depend on the formation of more invariant or abstract object representations or increased connections across the processing hierarchy that enable informed integration of incoming spatial features.

## Materials and Methods

### Participants

Behavioural participants were 74 children aged 3 - 13 years and 14 adults: 31 4- to 5-year-olds (mean age = 4.8, SD = 0.5 years, range = 3.1 - 5.7 years), 23 7- to 9-year-olds (mean age = 8.5, SD = 0.9 years, range = 6.8 - 9.7 years), 18 10- to 12-year-olds (mean age = 11.0, SD = 0.8 years, range = 10.2 - 13.7) and 13 adults (mean age = 22.2, SD = 2.2 years, range = 19.3 - 25.8 years). Age groups and sample sizes were defined based on effect sizes found in previous studies (Dekker et al. 2011, Yoon et al., 2007). Three 4- to 5-year-olds, one 7- to 9-year-old and one adult participant were excluded due to not completing the experiment or not complying with the task. An additional nine adult participants (mean age = 29.7, SD = 5.2 years, range = 23.7 - 38.9 years) completed piloting of the second stimuli set (‘trained’ set, see *Convolutional neural networks)*.

All participants had normal or corrected-to-normal vision, no reported history of eye disease, developmental or neurological disorder, or premature birth. Adults were recruited through the UCL Psychology Subject Pool (“SONA”), and received £10.50/hour compensation. Children were recruited through the UCL Child Vision Lab volunteer database, and received certificates, small toys, and transportation costs. Informed written consent was obtained from all adults and parents, and children gave verbal assent. The research was carried out in accordance with the tenets of the Declaration of Helsinki, and was approved by the UCL Ethics Committee (#1690/005).

### Apparatus

Stimuli were presented on a 22” touchscreen monitor (Iiyama ProLite T2252MTS 22”, 1900x1080 pixel resolution) driven by a MacbookPro, running Matlab R2015b with the Psychophysics Toolbox (Brainard, 1997).

### Behavioural Stimuli

Stimuli consisted of 20 two-tones created by processing greyscale photographs of objects, animals and faces in Matlab R2015b (MathWorks, Natick, MA, USA); images were first smoothed with a Gaussian filter and then thresholded to binarise pixel luminance to black or white to create images with obscured edges. Smoothing levels and binarisation thresholds varied per image. Images were selected from an original set of 41 stimuli following piloting with 13 adults and 16 children (4-10 years) for their effectiveness as two-tones, feasibility for testing young children, and to cover a range of difficulties for two-tone recognition. Selection criteria included high recognition accuracy (>95%) of greyscale versions at all ages, low recognition accuracy of naively viewed two-tones, and comparatively high recognition accuracy of cued two-tones. An additional 7 two-tones were generated without smoothing, creating easily recognisable two-tones, of which 3 were used in practice trials and 4 were used in ‘Catch’ trials, included to promote motivation, and obtain an index of attentiveness and task comprehension. All images were resized to a fixed height of 273.9 mm (680 pixels, ∼30 dva) on a mid-grey background. Text task prompts were displayed above the image and read aloud by the researcher.

### Procedure

Participants sat ∼30 cm in front of the touchscreen monitor that was in an upright position. Ambient lighting was controlled to maintain constant dim lighting. The experimenter was present throughout the procedure, and caregivers were sometimes present but did not engage with the procedure. Following task instruction, participants completed a training task in which a greyscale image was displayed and transformed gradually into a two-tone image that was unsmoothed and easy to recognise for all ages. To confirm that participants had understood that image content was maintained across the greyscale and two-tone, they were asked to point out corresponding features between the original greyscale image and the two-tone image when displayed side by side. Following this, participants completed 3 practice trials with unsmoothed two-tones. This was followed by 20 experimental trials with two-tones of varying difficulty, presented in a randomised order and interspersed with easy ‘catch’ trials every 5th trial, following which prize tokens were awarded to maintain motivation and a short break was allowed. Motivational, but uninformative, feedback was given after each trial to all participants.

Trials consisted of 3 sequential stages (Figure 1A). First, participants were permitted to free-view the uncued two-tone for an unlimited time, and asked to identify its content (stage 1). Once the participant answered, or was unwilling to guess after being prompted to, this two-tone was overlaid with its original greyscale image, preserving image size and position, and participants were asked to identify this image (stage 2). To demonstrate how the greyscale image is ‘transformed’ to the previously seen two-tone version, these images were then ‘morphed’ into the previously seen two-tone version via sequentially thresholding the greyscale with a decreasing number of channels represented (5 steps in total from 255 to 2 channels, with smoothing applied on the final step; 3-second duration in total). Participants were asked to confirm recognition of the two-tone by pointing out two defining features on the image (regions of interest, [ROIs]; stage 3). After completing all 20 experimental trials and 4 Catch trials, participants were then asked to locate the ROIs they had pointed out on the two-tone on each of the 25 corresponding greyscales. Verbal responses and screen coordinates of touched image locations were recorded for each trial. All task instructions were displayed as text above the stimulus image and also given verbally by the experimenter. The task duration was approximately 15 minutes including short breaks.

### Scoring of behavioural performance

To quantify image recognition, we measured ‘naive naming accuracy’, ‘cued pointing accuracy’, and ‘cued pointing distance’. *Naive naming accuracy*: Image names were scored as correct if the content was correctly identified at the basic category level (Rosch et al., 1976) - superordinate categories names (e.g., naming a ‘cow’ an ‘animal’) were scored as incorrect. Similar basic level or subordinate categories were accepted (e.g., a ‘tiger’ named as ‘cat’, or ‘scissors’ named as ‘shears’) as long as there was consistency in naming across the two-tone and greyscale (for all answers and coding scheme details per image, see Supplementary Table 1). Trials were excluded from Cued Pointing analyses if both targets were not attempted for both greyscale and two-tone conditions, and if participants were unable to accurately name the image content or locate each target in the greyscale condition. Cued pointing accuracy: pointed out feature location was scored as correct if the touched coordinates fell within a researcher-defined ROI demarcating the location of each target on both the two-tone and greyscale images. Cued pointing distance: the distance between each touched location for greyscale targets and the corresponding two-tone targets in millimetres. For the analysis of local errors, we considered eight targets for which incorrect features matching the target’s shape properties were present in the two-tone. Of the total errors made for these targets, the proportion whose touched coordinates fell within an ROI demarcating the incorrect shape-matching feature was counted for each age group. ROIs were equal in area to those used for the Pointing Accuracy measure described above.

### Convolutional neural networks

Pretrained AlexNet and NASNet-large models were acquired from and run with Matlab 2020a’s Deep Learning Toolbox (MathWorks, 2020). CORnet models (RT, Z and S) were acquired and run with the Python toolbox THINGSvision (Muttenthaler and Hebart, 2021). All models were run on a 2015 MacBook Pro. Due to poorer greyscale performance, CORnet-RT and CORnet-Z were not included in the results. All models were pre-trained to classify images into 1000 classes on the ImageNet dataset (Deng et al., 2009). Following the same coding scheme for human participants described above, CNN Classification Accuracy was determined separately for greyscale and two-tone images of ‘trained’ and ‘novel’ image sets (see *CNN Stimuli*) by whether the first-choice (highest weighted) classification was correct (with at least a basic-level match). Classification Probability was measured as the probability weighting of the highest weighted label scored as correct as per the coding scheme within the top 100 predictions.

### CNN Stimuli

Thirty-five images were selected from ImageNet to create additional greyscales and two-tones, chosen due to similarities with the behavioural stimuli set and suitability for two-tone transformation. Resulting greyscales and two-tones were dynamically cropped to a 680*680-pixel square to minimise loss of image content. Image piloting was carried out in eight additional adult participants using an adapted version of the behavioural task described above: Images were shown in 7 blocks of 5 images, where 5 naive two-tone trials were followed by 5 corresponding greyscale trials and finally 5 cued two-tone trials, with image order randomised within conditions. Recognition was determined via Naming Accuracy (as above) for all three conditions, with an additional perceptual check (‘Which way is the animal/object facing?’) asked as confirmation for scoring cued trials correct. Following piloting, 20 images were selected to match the behavioural stimuli on smoothing, thresholding and adult performance levels. Of the 20 behavioural stimuli described above, 19 that were not found within the ImageNet dataset (one greyscale - two-tone pair excluded; behavioural image 6, see Supplementary Table 1) comprised the ‘novel’ image set. Images of both sets were resized and triplicated across RGB channels to match model input sizes (AlexNet: 227*227*3 pixels, NASNet: 331*331*3 pixels, CORnet-S: 224*224*3 pixels).

### Statistical analyses

To analyse responses, we used a mixed-effects modelling approach, as this allowed us to account for different numbers of trials per condition and age groups, and random effects of individual participants and images. We used generalised linear models estimated using the lme4 library in R (Bates et al., 2015). To compare naming we used multilevel logistic regression models with crossed random intercepts for participants and images. To compare pointing accuracy, we used multilevel logistic regression models with crossed random intercepts for participants, images, and ROI area in pixels. For distance measures, we used a multilevel generalised linear model with a log link function (as pointing error is strictly positive, distance data were log-transformed to ensure normality), with crossed random intercepts for participants, images, and ROI area. To test for age trends, we tested each model against a reduced model without the age effect of interest, using a likelihood ratio test (data and analysis code available on request). We performed Wald tests on the coefficients for pair-wise age group comparisons.

## Acknowledgments

The research was supported by grants from the National Institute for Health Research (NIHR) Biomedical Research Centre (BRC) at Moorfields Eye Hospital NHS Foundation Trust and UCL Institute of Ophthalmology, the Economic and Social Research Council (ESRC) of the UKRI (#ES/N000838/1); and Moorfields Eye Charity (R160035A, R190029A, R180004A).

