## Supplementary Materials for "Perceptual reorganisation from prior knowledge emerges late in childhood"

**Supplementary Table 1**: Naming Accuracy Coding Scheme

Table shows accepted answers for Catch trials (rows 1-4), Behavioural Stimuli (rows 5-24) and 'Trained' Stimuli (rows 25 - 45). Asterisk denotes answers which were only accepted if consistent with corresponding greyscale answers (for behavioural trials) or if no other more applicable classification was available (for CNNs).

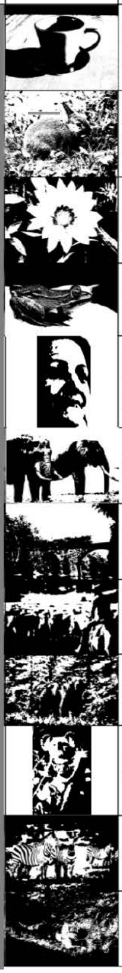

| | Greyscale | Two-tone | Accepted Answers |
|---|---|---|---|
| Catch 1 | | | Cup, Mug, Coffee Mug, Teacup, Glass*, Teapot*, Coffeepot* |
| Catch 2 | | | Rabbit, Type of rabbit, Bunny, Hare Squirrel*, Badger* |
| Catch 3 | | | Flower, Sunflower, Daisy |
| Catch 4 | | | Frog, Type of frog, Toad, Lizard*, Type of lizard* |
| Behav. 1 | | | Woman, Lady, Girl, Face, Person, Human, Man*, Eyes*, Bob Marley* |
| Behav. 2 | | | Elephant, Elephants, Tusker |
| Behav. 3 | | | Train, Bridge, Type of bridge |
| Behav. 4 | | | Cowboy, Horseman, Man/Person on horse, Horse, Person*, Man*, Cow*, Cowboy hat* |
| Behav. 5 | | | Bear, Brown bear, Polar Bear*, Panda*, Bears* |
| Behav. 6 | | | Koala, Sloth*, Monkey*, Bear*, Squirrel* |
| Behav. 7 | | | Zebra, Zebras |
| Behav. 8 | | | Tiger, Lion*, Cat*, Wildcat* |

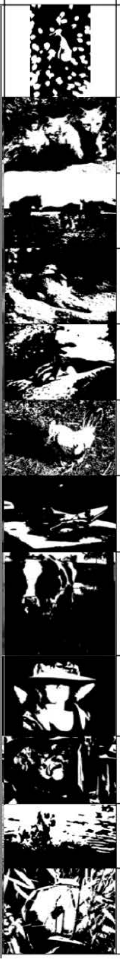

| | Greyscale | Two-tone | Accepted Answers |
|---|---|---|---|
| Behav. 9 | | | Fish, Nemo, Anemone fish, Coral reef |
| Behav. 10 | | | Fox, Foxes, Fox cubs, Type of fox, Cats*, Kittens*, Dogs*, Pups*, Wolves*, Cubs*, Hyena*, Skunk* |
| Behav. 11 | | | Horses, Horse, Cows*, Camel*, Yeehah animal*, Horse cart*, Water buffalo* |
| Behav. 12 | | | Alligator, Crocodile, Lizard* |
| Behav. 13 | | | Chipmunk, Squirrel, Rat, Mouse, Weasel |
| Behav. 14 | | | Rooster, Cockerel, Chicken, Hen, Bird, Cock, Duck* |
| Behav. 15 | | | Scissors, Knife*, Cutting* |
| Behav. 16 | | | Cow, Cows, Ox, Moo* |
| Behav. 17 | | | Woman, Lady, Girl, Person, Human, Face, Hat, Cowboy hat, Sombrero, Queen, Witch, Man*, Detective* |
| Behav. 18 | | | Man, Boy, Face, Person, Human, Grandpa, Old lady, Cowboy Hat*, People*, Witch*, Mouth* |
| Behav. 19 | | | Dog, Breed of dog, Puppy, Otter* |
| Behav. 20 | | | Panda, Giant panda, Bear, Koala* |

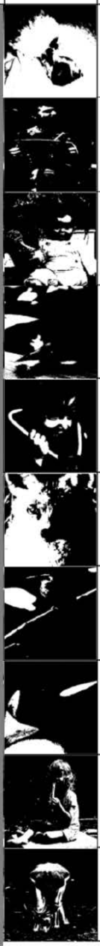

| | Greyscale | Two-tone | Accepted Answers |
|---|---|---|---|
| Trained 1 | | | Dog, Breed of dog |
| Trained 2 | | | Man, Human, Person, Accordion |
| Trained 3 | | | Baby, Toddler, Maraca |
| Trained 4 | | | Dog, Breed of dog |
| Trained 5 | | | Girl, Boy, Doctor, Stethosco pe |
| Trained 6 | | | Fox, Type of fox, Dog, Wolf, Type of Wolf |
| Trained 7 | | | Bird, Type of bird |
| Trained 8 | | | Penguin, King penguin, Bird, Type of bird |
| Trained 9 | | | Girl, Boy, Child, Ice lolly* |
| Trained 10 | | | Deer, Antelope, Gazelle, Ox |

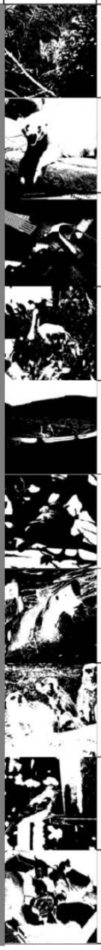

| | Greyscale | Two-tone | Accepted Answers |
|---|---|---|---|
| Trained 11 | | | Lynx, Cat, Big Cat, Snow Leopard |
| Trained 12 | | | Polar bear, Ice bear, Bear |
| Trained 13 | | | Hammer, Tools |
| Trained 14 | | | Bird, Type of bird |
| Trained 15 | | | Boat, Canoe, Paddle* |
| Trained 16 | | | Bird, Type of bird |
| Trained 17 | | | Shark, Great White, Whale, Killer Whale |
| Trained 18 | | | Meerkat, Meerkats, Dog*, Dogs*, Type of Dog* |
| Trained 19 | | | Bird, Type of bird |
| Trained 20 | | | Dog, Dogs, Puppies, Breed of dog |

**Supplementary Figure 2**: Catch Trial Performance

**A)** Naive naming accuracy of two-tones (coloured circles), and Catch images ('easy' two-tones created without smoothing; white circles). Small markers show participant means, large markers show age group means, error bars show bootstrapped 95% Confidence Intervals. **B)** Pointing accuracy for two-tones (coloured circles) and Catch trials (white circles) following greyscale exposure, as measured by percentage of touched locations falling within the predefined correct area for each target. Markers and error bars are as in Panel A. **C)** Catch trial performance of CNNs (greyscale bars) and participant age groups (coloured bars). Error bars show bootstrapped 95% Confidence Intervals for participant age groups.

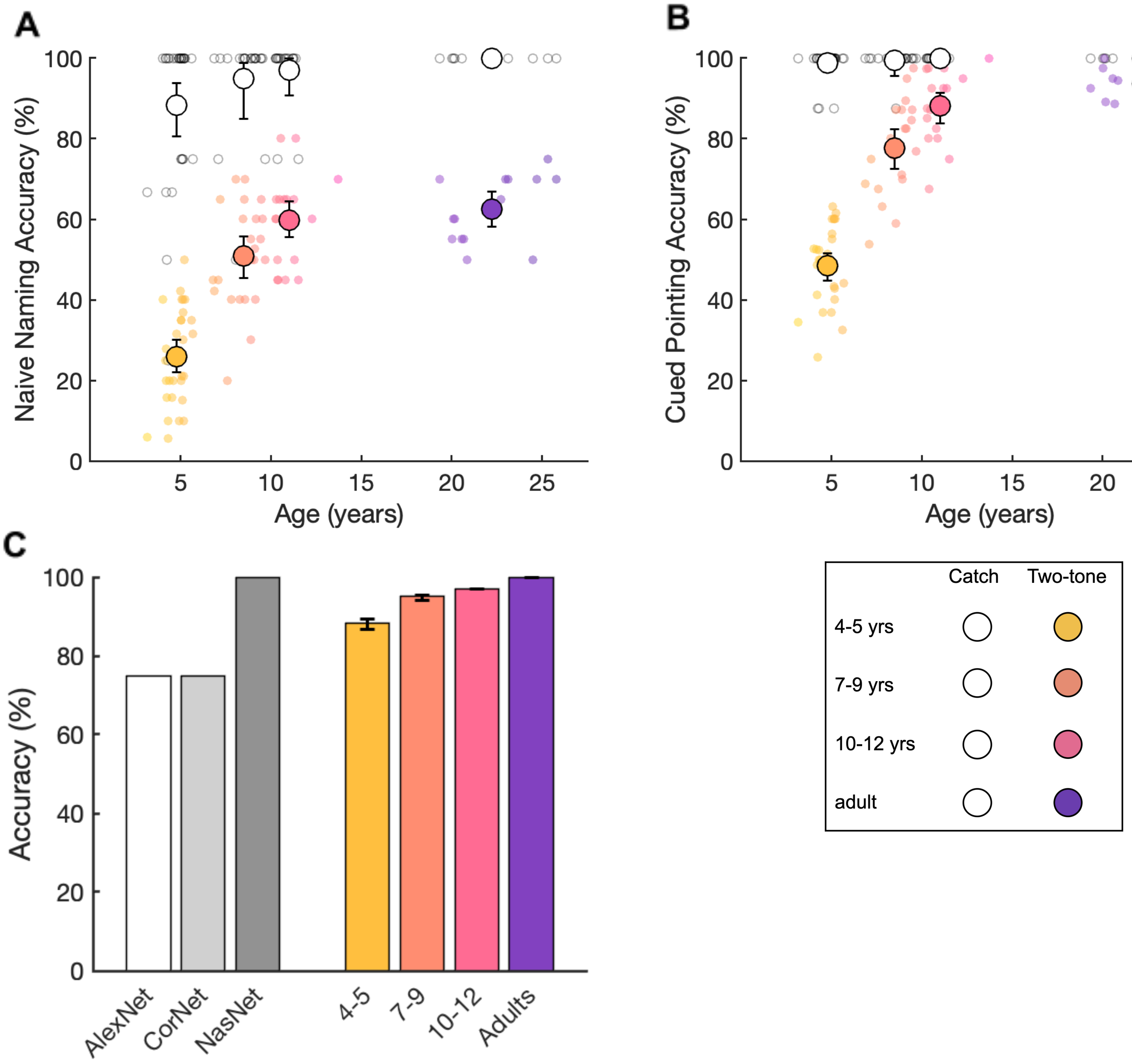

**Supplementary Figure 3**: Feature Localisation Performance per Image.

Catch (top row) and two-tone images additional to those shown in Figure 4 (rows 2-5) with 4- to 5-year-olds' (yellow markers) and adults' (purple markers) cued two-tone pointing: target features 1 and 2 are shown in subsequent images. Target prompts were (1) 'the cup's handle', (2) 'the cup's saucer', (3) 'the frog's eye', (4) 'the frog's leg', (5) 'the middle of the flower', (6) 'the edge of one of the petals', (7) 'the rabbit's eye', (8) 'the rabbit's ear', (9) 'the left elephant's' tusk', (10) 'the right elephant's mouth', (11) 'the front of the train', (12) 'the column of the bridge', (13) 'the bear's left ear', (14) 'the bear's nose', (15) 'the koala's nose', (16) 'the koala's hand', (17) 'the left zebra's eye', (18) 'the right zebra's back leg', (19) 'the tiger's left ear', (20) 'the tiger's tail', (21) 'the fish's eye', (22) 'the fish's side fin', (23) 'the left horse's ears', (24) 'the right horse's nose', (25) 'the alligator's nose', (26) 'the alligator's eye', (27) 'the chipmunk's eye', (28) 'the chipmunk's tail', (29) 'the chicken's beak', (30) 'the chicken's eye', (31) 'the scissors' bolt', (32) 'the scissors' right handle', (33) 'the cow's nose', (34) 'the cow's rear', (35) 'the lady's nose', (36) 'the lady's hat', (37) 'the man's ear', (38) 'the man's nose', (39) 'the dalmatian's left eye' and (40) 'the dalmatian's nose'.

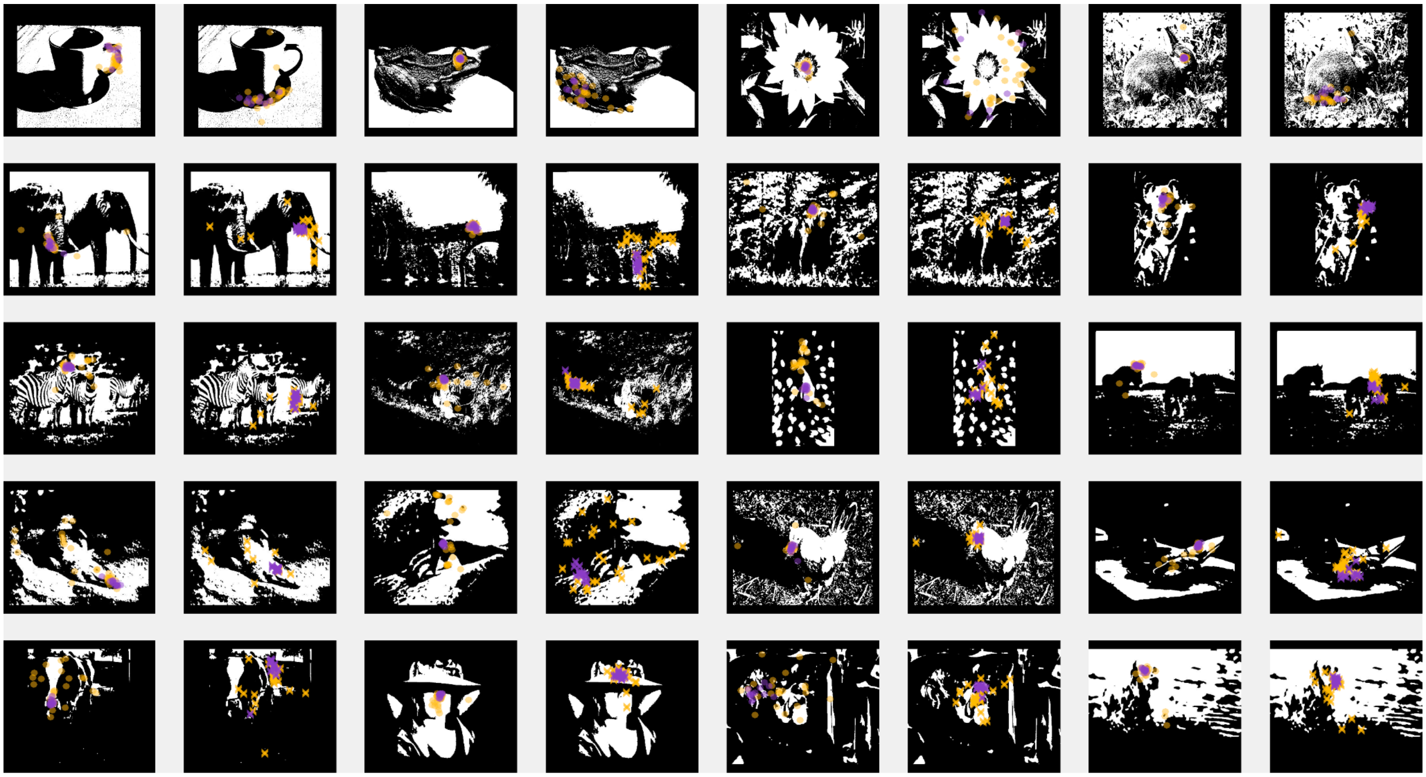

**Supplementary Figure 4:** Image-wise correlations of human and CNN performance across conditions.

Comparable results were found for Spearman Rank correlations.

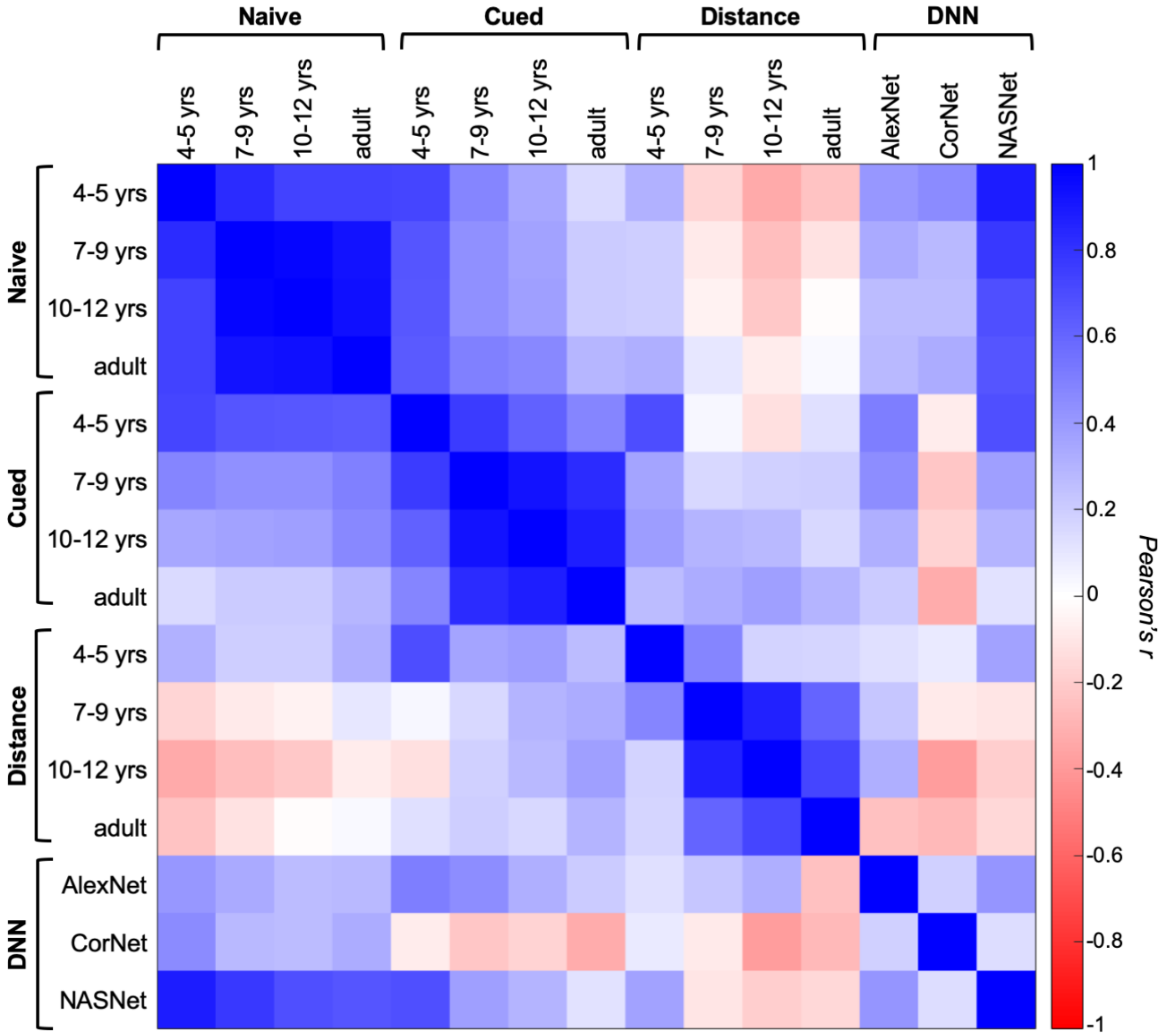

**Supplementary Materials 5: Computational model for computing natural scene statistics.**

Low-level spatial information in the image such as edge and contrast metrics convey information about image content. This information has been found to guide parsing of objects in complex visual scenes (Groen et al., 2018; Seijdel et al., 2020). In two-tone recognition, the disambiguation of obscured edges using object knowledge is thought to play an important role in the parsing process. We hypothesised that an image recognition strategy relying on spatial information about edges would be adversely affected by disruptions of this information following two-tone transformation. We tested this by quantifying several spatial image statistics for the greyscale and two-tone conditions of each image, as well as the change between these conditions. We predicted negative correlations between performance and the change in greyscale and two-tone image statistics, in particular for younger children who may rely more heavily on the image features rather than object knowledge.

Edge Density, the percentage of image pixels that represent edges, of the images in our dataset decreased following two-tone transformations. Larger reductions in Edge density imposed by converting from greyscale to two-tone correlated with poorer naive two-tone recognition in 4- to -5-year-olds' (see 'Δ Edge Density', Supplementary Table 5). An overall higher two-tone Edge Density correlated with better Cued Accuracy for this age group (see 'Edge Density', Supplementary Table 5).

We also computed biologically principled summary scene statistics using a gist extraction model that emulates how the magno- and parvocellular pathways process low-level features with neurally-plausible image filtering (Groen et al., 2013). This model first convolves the image with a range of local edge filters that mimic parvo and magnocellular spatial filters (Scholte et al., 2009), the responses of which were rectified and divisively normalised following the LGN suppressive field approach (Bonin et al., 2005). One filter per image location was selected using minimal reliable scale selection (Ghebreab et al., 2009), and the resulting spatially filtered images are then pooled into two summary statistics: Contrast Energy (CE) and Spatial Coherence (SC). To compute CE, the edge filter responses for the parvocellular pathway are averaged, resulting in a measure of mean image contrast, with high values reflecting more contrast. To compute Spatial Coherence (SC), the variability of edge filter responses is computed for the magnocellular pathway, resulting in a measure of contrast variability, with high values reflecting more contrast variations. In Groen et al's (2013) study, these statistics were computed across a limited visual field area, to account for the fact that participants are fixating centrally on a briefly (100 ms) presented stimulus. However, in our study, images were free-viewed for extended periods, so we averaged statistics across the entire image (i.e., a 15-degree radius). Groen et al. (2013) showed that in adults, these model outputs predict perceived naturalness of the image and correlate with the EEG signal amplitude recorded across the occipital cortex whilst the image is categorised. We recently showed that this is also the case in children aged 6 years and upward, although less variance in the occipital EEG was explained by the model in the youngest children (Chow-Wing-Bom et al., 2019).

Here, the conversion of greyscale images to two-tones was found to increase CE and reduce SC overall (see Supplementary 6). Greater alterations of these parameters corresponded with poorer Naive and Cued two-tone recognition in 4- to 5-year-olds, but no other age group (see Supplementary 7: ‘Δ CE’ and ‘Δ SC’). These data are tentatively in line with age-related reductions in the reliance on intact image statistics when object knowledge is available.

**Supplementary Table 5**: Pearson’s correlations of image properties and performance by age group.

Image properties are (1) Thresholding and (2) Smoothing levels used for the two-tone transformation of each image, (3) Edge Density, (4) Contrast Energy and (5) Spatial Coherence of resulting two-tone images, and Change between greyscale and two-tone images (Δ) for (6) Edge Density, (7) Contrast Energy and (8) Spatial Coherence. Correlations are uncorrected for multiple comparisons, and comparable results were found for Spearman Rank correlations.

| | Naïve Naming Accuracy | | | | | | | Cued Pointing Accuracy | | | |
|---|---|---|---|---|---|---|---|---|---|---|---|
| | 4-5 yrs | 7-9 yrs | 10-12 yrs | Adults | AlexNet | CorNet-S | NASNet | 4-5 yrs | 7-9 yrs | 10-12 yrs | Adults |
| **Threshold** Pearsons’ *r* | **-0.529** | -0.384 | -0.293 | **-0.410** | -0.036 | -0.251 | **-0.469** | -0.300 | -0.306 | -0.281 | -0.055 |
| *p* | **0.007**** | 0.064 | 0.164 | **0.048*** | 0.899 | 0.387 | **0.024*** | 0.154 | 0.145 | 0.183 | 0.797 |
| **Smoothing** Pearsons’ *r* | **-0.695** | **-0.673** | **-0.611** | **-0.638** | -0.498 | -0.101 | **-0.625** | **-0.448** | **-0.442** | -0.381 | -0.162 |
| *p* | **0.0002***** | **0.0003***** | **0.001**** | **0.0008***** | 0.059 | 0.731 | **0.001**** | **0.028*** | **0.031*** | 0.066 | 0.450 |
| **Edge Density** Pearsons’ *r* | 0.389 | 0.382 | 0.360 | 0.371 | 0.414 | 0.052 | **0.467** | **0.467** | 0.316 | 0.275 | 0.185 |
| *p* | 0.060 | 0.065 | 0.084 | 0.075 | 0.125 | 0.861 | **0.025*** | **0.021*** | 0.133 | 0.193 | 0.388 |
| **Contrast Energy** Pearsons’ *r* | **0.525** | 0.317 | 0.249 | 0.343 | -0.127 | 0.387 | **0.442** | **0.417** | 0.275 | 0.314 | 0.160 |
| *p* | **0.008**** | 0.131 | 0.240 | 0.100 | 0.653 | 0.172 | **0.035*** | **0.043*** | 0.193 | 0.135 | 0.456 |
| **Spatial Coherence** Pearsons’ *r* | 0.283 | 0.247 | 0.234 | 0.322 | -0.031 | 0.200 | 0.350 | 0.306 | 0.108 | 0.113 | 0.005 |
| *p* | 0.180 | 0.245 | 0.271 | 0.125 | 0.911 | 0.493 | 0.102 | 0.146 | 0.616 | 0.599 | 0.982 |
| **Δ Edge Density** Pearsons’ *r* | **0.525** | 0.372 | 0.261 | 0.197 | 0.498 | 0.110 | **0.538** | 0.298 | 0.214 | 0.158 | 0.137 |
| *p* | **0.008**** | 0.073 | 0.219 | 0.356 | 0.059 | 0.708 | **0.008**** | 0.157 | 0.315 | 0.460 | 0.524 |
| **Δ Contrast Energy** Pearsons’ *r* | **0.6385** | 0.390 | 0.311 | 0.398 | 0.008 | 0.475 | **0.491** | **0.448** | 0.300 | 0.283 | 0.143 |
| *p* | **0.0008***** | 0.060 | 0.140 | 0.054 | 0.977 | 0.086 | **0.017*** | **0.028*** | 0.155 | 0.181 | 0.506 |
| **Δ Spatial Coherence** Pearsons’ *r* | **-0.454** | -0.376 | -0.329 | -0.352 | -0.367 | -0.161 | **-0.581** | -0.293 | -0.015 | 0.090 | 0.101 |
| *p* | **0.026*** | 0.070 | 0.117 | 0.091 | 0.178 | 0.583 | **0.004**** | 0.164 | 0.946 | 0.675 | 0.639 |

**Supplementary Figure 5:** Two-tone and greyscale differences in image statistics.
**A)** Contrast Energy and Spatial Coherence of greyscale (blue borders), two-tone (pink borders) and catch (yellow borders) images. **B)** Comparison of average Edge Density, Contrast Energy and Spatial Coherence for greyscale, catch and two-tone stimuli. Error bars show bootstrapped 95% Confidence Intervals.

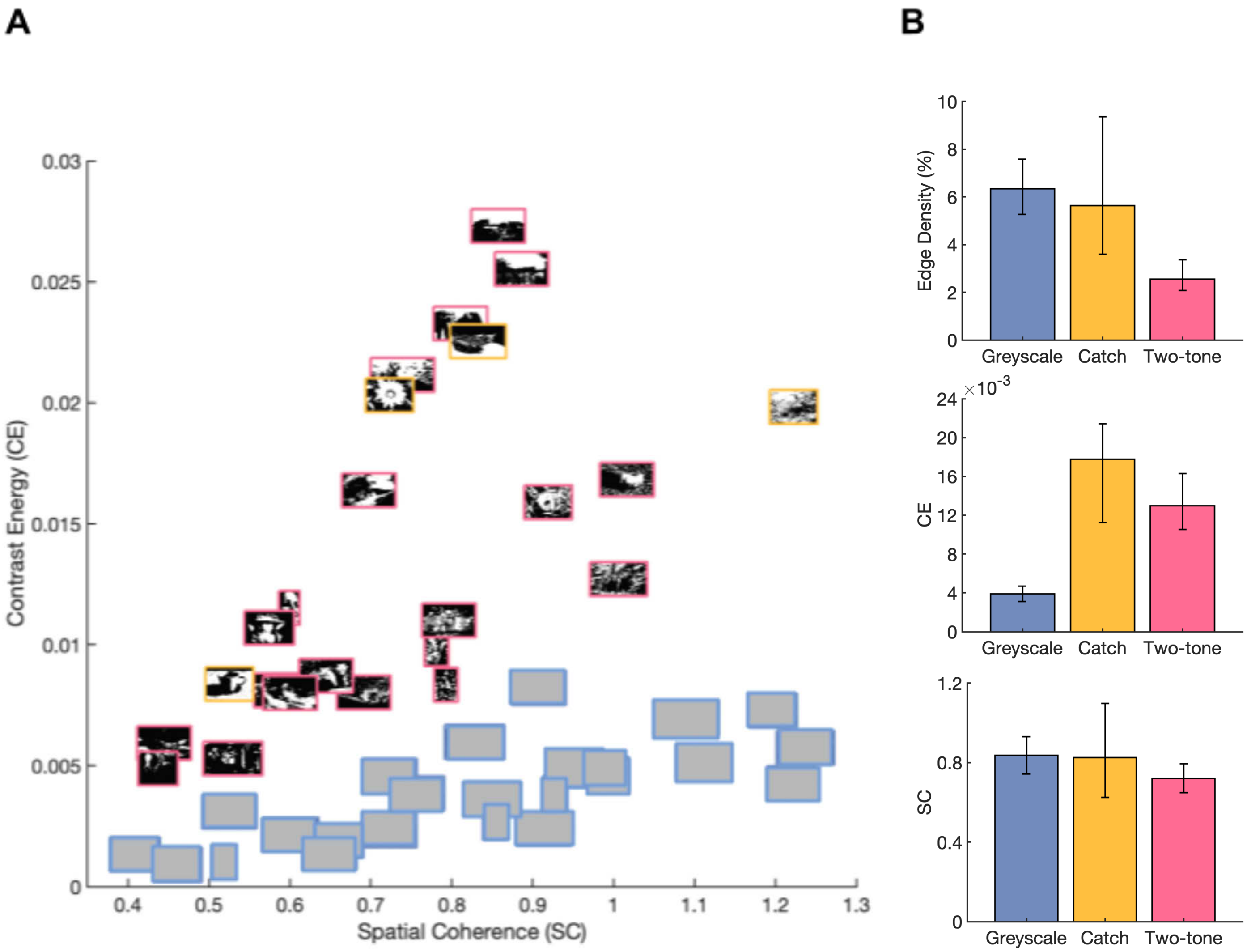
